# The dynamic expressome of *Pseudomonas putida*: condition-dependent remodelling of the RNA polymerase interactome

**DOI:** 10.64898/2026.11.17.767251

**Authors:** Borja Pitarch, Florencio Pazos, Víctor de Lorenzo, Huseyin Tas

## Abstract

As in many bacteria, the α_2_ββ′ω core RNA polymerase (RNAP) of *Pseudomonas putida* interacts *in vivo* with sigma factors, transcriptional regulators and auxiliary proteins whose relative abundance varies with the physiological state. Here, we used an *in vivo* immunocapture approach to examine condition-dependent changes in the RNAP interactome of *P. putida* KT2440. His-, Myc- and E-tags were separately introduced at the C-terminus of RpoC (β′), a permissive, surface-exposed site accessible to cognate antibodies. Myc-tagged RNAP complexes were immunocaptured with magnetic beads and analyzed by LC-ESI-MS/MS and computational methods. The results revealed condition-dependent association of the RNAP core (RpoA, RpoB, RpoC and RpoZ) with distinct pools of sigma factors, as well as ribosomal proteins, RNA-processing factors and metabolic enzymes. Nutrient starvation, nutrient-rich growth and high temperature altered the composition and relative representation of RNAP-associated proteins. Despite the limited physiological conditions examined, our results support a model in which RNAP acts as a dynamic organizational hub of the bacterial expressome, integrating transcription with translation, RNA maturation and central metabolism.

## 1. INTRODUCTION

RNA polymerase (RNAP) is the central catalytic machinery of gene expression in all living systems. In bacteria, the housekeeping RNAP core enzyme is composed of five subunits arranged as α_2_ββ′ω, which associate with alternative sigma (σ) factors to form transcriptionally competent holoenzymes capable of recognizing specific promoter sequences and initiating transcription (Murakami and Darst, 2003; Feklístov *et al*., 2014). The molecular basis of transcription initiation has been extensively characterized in model organisms such *as Escherichia coli* and *Bacillus subtilis*, where purified RNAP, sigma factors, transcriptional regulators, and DNA templates have enabled the faithful reconstitution of many aspects of promoter-dependent transcription *in vitro* (Haldenwang, 1995; Saecker *et al*., 2011; Helmann, 2019). However, the operation of RNAP *in vivo* occurs within a densely crowded and highly structured cytoplasmic environment, where the transcriptional machinery is continuously exposed to a multitude of potential protein partners that may influence its activity, stability, localization, and regulatory properties (Jin *et al*., 2013; Muskhelishvili *et al*., 2021).

Increasing evidence suggests that RNAP does not function as an isolated molecular machine but rather as the central component of a dynamic network of protein–protein interactions. Beyond its well-established associations with sigma factors, transcriptional activators and repressors, elongation factors, and termination proteins, RNAP participates in additional interactions that functionally connect transcription with other cellular processes, including translation, metabolic and stress responses, and chromosome organization (Jin *et al*., 2013; Browning and Busby, 2016; Blaha and Wade, 2022). Such RNAP-centered interactomes have been investigated in several bacterial species, using affinity purification or immunoprecipitation of RNAP components coupled to mass spectrometry to identify proteins associated with the transcriptional machinery (Lee *et al*., 2008; Pelletier *et al*., 2008; Kriel *et al*., 2020; Vasilyev *et al*., 2024), revealing a surprisingly complex molecular entourage in its surrounding. Yet, the composition and dynamics of these interactomes remain poorly understood, particularly in environmentally relevant microorganisms and under physiologically meaningful conditions. Among emerging bacterial platforms for biotechnology, *Pseudomonas putida* has attracted considerable attention because of its exceptional metabolic versatility, robustness, and tolerance to environmental stresses (Nikel and de Lorenzo, 2018). In particular, the strain *P. putida* KT2440 have become widely adopted as chassis strains for industrial biotechnology, synthetic Biology (Tas, 2020; Tas *et al*., 2021) and environmental applications (de Lorenzo *et al*., 2024). Their ability to withstand toxic chemicals, oxidative stress, solvent exposure, and other harsh conditions distinguishes *P. putida* KT2440 from traditional microbial workhorses and has motivated extensive efforts to exploit these traits for the production of chemicals, materials, and bioremediation agents (Nikel and de Lorenzo, 2018).

The core transcriptional machinery of *P. putida* shares many features with that of *E. coli*, as illustrated by the ability of numerous promoters to function across both species and, in several cases, to initiate transcription from equivalent sites (Bagdasarian *et al*., 1983; Pinkney *et al*., 1987; Sawers *et al*., 1989; Roldán and Amarelle, 2024). Nevertheless, promoter activities are often not quantitatively transferable, they show contextual dependencies (Tas *et al*., 2021) indicating that the molecular context in which transcription occurs differs substantially between the two organisms. Such differences may arise from variations in sigma factor repertoires, transcriptional regulators, chromosome architecture, metabolic state, and the composition of the RNAP interactome itself. Despite the growing importance of *P. putida* as a model chassis, comparatively little is known about the proteins that physically associate with its RNAP or how these associations change in response to environmental stimuli.

On this background, we wondered whether the remarkable physiological robustness of *P. putida* is reflected in distinctive features of its transcriptional machinery and, in particular, in the dynamic network of proteins that accompany RNAP during gene expression. To systematically characterize the *P. putida* RNAP interactome under physiological conditions, we adopted an *in vivo* immunocapture strategy based on chromosomal epitope tagging of RpoC, the β′ subunit of RNAP. This approach builds on previous studies showing that genetically introduced affinity tags at the permissive C-terminal region of RpoC can provide an effective handle for purification of intact RNAP together with associated proteins, while preserving enzyme functionality (Lee *et al*., 2008; Svetlov and Artsimovitch, 2015). Immunoprecipitation then enabled the isolation of RNAP-containing complexes directly from extracts of living cells and the subsequent identification of associated proteins. Using this methodology, we examined the composition of RNAP-associated protein assemblies under different physiological conditions, including heat-shock stress. The resulting datasets revealed not only the expected interactions between RNAP and components of the transcriptional machinery, such as sigma factors and termination or antitermination proteins, but also a diverse collection of additional partners, including ribosomal proteins, metabolic enzymes, and proteins of previously unknown connection to transcription. Comparative analysis of the recovered complexes further demonstrated that the RNAP interactome is highly dynamic and responsive to environmental conditions. More broadly, the results support a view of bacterial RNA polymerase not merely as a DNA-dependent enzyme but as the hub of a condition-dependent molecular network (the *expressome*; Kohler *et al*., 2017) that coordinates gene expression with the metabolic and physiological state of the cell.

## 2. MATERIALS AND METHODS

### 2.1 Strains, culture conditions and general procedures

Unless otherwise indicated, *Pseudomonas putida* KT2440 and its derivatives were grown in Luria– Bertani (LB) medium at 30°C. Liquid cultures in flasks were incubated with aeration in an orbital shaker. Growth experiments in 96-well microtiter plates were performed using a SpectraMax M2 microplate reader (Molecular Devices, Sunnyvale, CA, USA). For heat-shock experiments, cultures were grown under the same conditions except that the incubation temperature was shifted to 42°C for 3 h. By default, cultures were initiated from fresh colonies and grown overnight to saturation under the conditions described above. These overnight cultures were subsequently used to inoculate fresh medium to obtain synchronized second-generation cultures for the experiments. For determination of doubling times, three independent experiments were performed using 20 ml LB medium in 100-ml flasks incubated at 30°C with aeration. Cultures were inoculated from overnight cultures to an initial OD_600_ of 0.02, and growth was monitored over time. All DNA constructs were verified by Sanger sequencing using standard procedures. Primers used for construct generation and sequence verification are listed in Supplementary Table S1.

### 2.2. Immuno-genetic tagging of RpoC (β’ subunit of RNAP)

Three epitope tags for immunodetection/capture were separately introduced to the sequence of the structurally permissive C-terminus of the *rpoC* gene of *P. putida* (Fig. 1A). These included the His-tag (10xHis), the c-Myc-tag (EQKLISEEDL) and the E-tag (GAPVPYPDPLEPR) (Terpe, 2003). To this end, the corresponding DNA sequences were inserted prior to the stop codon TAA of the native gene, preceded either by two (for His-tag) or 4 (for Myc-tag and E-tag) Gly codons for improving the accommodation and exposure of the labels on the protein surface (Fig. 1B). The thereby engineered sequences were assembled in vector pEMG *and* delivered to the chromosomally borne *rpoC* gene through homologous recombination, following the protocol of (Martínez-García and de Lorenzo, 2011; Martínez-García and de Lorenzo, 2012). A summary of the construction flow is sketched in Supplementary Fig. S1. As explained in the text and Supplementary Table S2, these procedures resulted in strains *P. putida* TAS108 (β′-10xHis), *P. putida* TAS110 (β′-Myc) *and P. putida* TAS114 (β′-E-tag). Additional strains bearing a fusion of the reference promoter PBG37 promoter to a GFP reporter (Zobel *et al*., 2015) were derived from TAS108 and TAS110, giving rise to TAS103 (*P. putida* KT2440 *rpoC*::HisTag *att*Tn*7*::PBG37-msfGFP) and TAS106 (*P. putida* KT2440 *rpoC*::MycTag *att*Tn*7*::PBG37-msfGFP), respectively.

**Figure 1.**
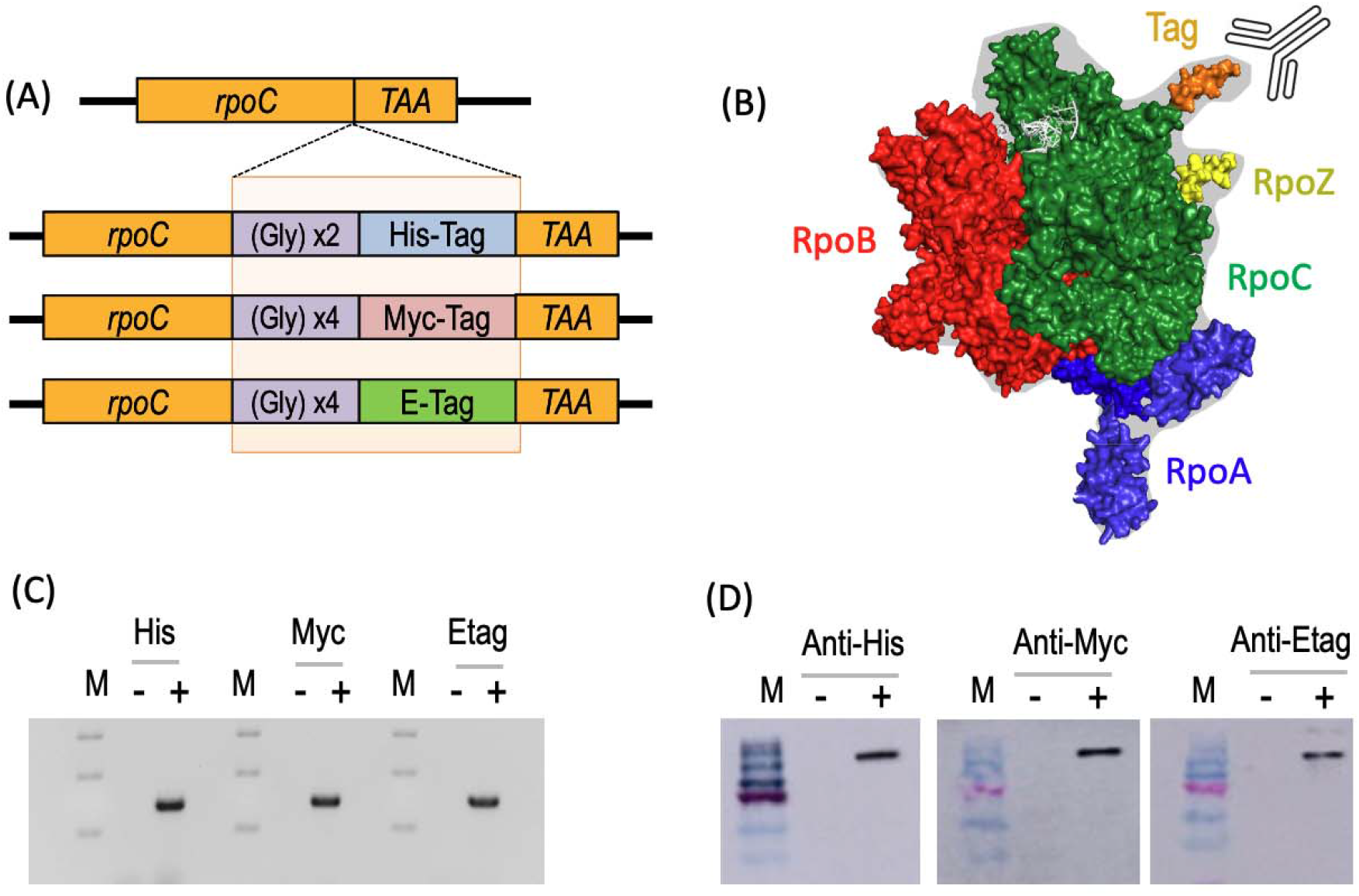
Labelling RNA of *P. putida* KT2440 with permissive immuno-tags. **(A)** Addition of 3 immunogenic epitopes sequences downstream of the *rpoC* gene before the stop codon. As indicated, the tags entered in the genome of the strain include Hisx6, c-Myc tag and E-tag. Gly residues were added between the RpoC and the tag as indicated in order to give flexibility and exposure new epitope (not to scale). RpoC is ∼155 kDa and each tag is ∼1 kDa. **(B)** Three-dimensional model of the RNAP of *P. putida*. The structure was derived from E. coli ‘s RNAP (PDB ID: 8PBL) whose alignment gives an excellent root mean square deviation of atomic positions (RMSD) of 1.456. The image shows the spatial arrangement of each of the units and the location of the immunogenic tag at the C-terminus of the β’ subunit (RpoC). **(C)** PCR verification of the labels inserted in *rpoC* by using the tag-specific primer sets indicated in Supplementary Table S1. M, size markers. - lanes, non-modified wild type controls. +, modified strains. Tagged strains show the right size of the amplification product (∼0.7 kB). (D) Western Blot analysis for verifying each labelled strain with its corresponding tags. The band sizes (∼155 kDa) correspond to RpoC size.

### 2.3 Protein extraction, RNAP pull-down and verification

Protein complexes were extracted using an adapted BugBuster Protein Extraction Reagent/Lysonase Bioprocessing Reagent protocol (Novagen, Merck). Overnight second-generation cultures were prepared by diluting primary overnight cultures 1:1.000 into fresh LB medium and incubating at 30 °C with shaking up mid-exponential phase. The OD_600_ of the resulting cultures was measured and an amount of bacteria corresponding to an OD_600_ ∼ 2 was collected by centrifugation (13.000 rpm, 30 min, room temperature). Cell pellets were resuspended in 1 mL BugBuster Protein Extraction Reagent (Merck Millipore) supplemented with 1 µL Lysonase Bioprocessing Reagent (a mixture of recombinant lysozyme and nucleases) following provider’s instructions. Suspensions were incubated at room temperature with gentle shaking (125 rpm) for 10 min, during which progressive clarification of the suspension was observed. Cell debris was subsequently removed by centrifugation (14,000 rpm, 60 min, 4 °C), and the cleansed supernatant containing soluble protein complexes was transferred to a fresh tube for immediate processing or stored at −80 °C. For heat-shock experiments, cells were grown at 30 °C to exponential phase OD_600_ ∼ 0.4), shifted to 42 °C for 3 h (at which point they stopped growing) and subsequently harvested and processed as before. Protein complexes associated with c-Myc-tagged RNA polymerase (RNAP) were isolated from clarified lysates by incubation with 50 µL of anti-c-Myc MicroBeads for 30 min on ice and further separation in μColumns placed in a μMACS magnetic separator (Miltenyi Biotec). After washing according to the manufacturer’s protocol, retained RNAP-containing complexes were recovered using elution buffer preheated to 95 °C. The resulting immunoprecipitates were used for downstream analyses, including SDS-PAGE, Western blotting and proteomic characterization.

### 2.4. Western blot analysis

Western blotting was used to verify the presence of the epitope-tagged proteins in the corresponding samples and immunoprecipitates. Following SDS-PAGE (Laemmli, 1970) the stacking portion of the gel (4% polyacrylamide) was removed and proteins in the resolving 8% gel were transferred onto a membrane. Prior to transfer, the membrane was activated by immersion in methanol for 1 min and subsequently equilibrated in transfer buffer (25 mM Tris, 192 mM glycine, pH 8.3, 20% v/v methanol). The gel and activated membrane were assembled between transfer buffer-soaked filter sponges, taking care to eliminate air bubbles between the different layers. The assembled sandwich was then immersed in transfer buffer and the setup run with a constant current of 0.1 A for 30 min. Following transfer, the membrane was incubated for 1 h at room temperature in buffer A (0.1 % Tween 20 in PBS: 8 mM Na2HPO4, 1.5 mM KH2PO4, 3 mM KCl, 137 mM NaCl, pH 7.0) containing 3% (w/v) skim milk to block non-specific protein-binding sites. The membrane was subsequently incubated for 1 h at room temperature with the peroxidase (POD)-conjugated antibody indicated in each case diluted 1:10.000 in the same blocking solution. After antibody incubation, the membrane was washed four times for 5 min each with Buffer A under gentle agitation to remove unbound antibody. Antibody-bound proteins were detected using a peroxidase-compatible chemiluminescent substrate. The membrane was incubated with the freshly prepared detection reagent for 1 min in the dark and immediately subjected to imaging with an Amersham™ Imager 600. Chemiluminescent signals were recorded using appropriate exposure times to visualize the protein bands of interest.

### 2.5. Proteomic analyses

The pools of proteins pulled down under the various conditions were subsequently investigated with ultra-short gradient Liquid Chromatography coupled with Electrospray Ionization Tandem Mass Spectrometry (LC-ESI-MS/MS) as explained in (Arcos *et al*., 2014). and (Bodega *et al*., 2018). Proteomics results provided a Table with the abundances of proteins in all four different conditions. Normalised abundances were calculated dividing the abundance of an experimental condition by the abundance of the control condition (S1, proteins pulled down by the anti c-Myc antibody in an extract of a wilt-type strain with a non-tagged RNAP, see below). A threshold of 10 in the normalised abundance is imposed to determine that a protein is enriched in the condition. Having the list of enriched proteins for each condition, we checked at STRING database (Szklarczyk *et al*., 2019; https://string-db.org/ , version 11.5) which of these proteins had a described interaction with RpoC and have a confidence score above 0.900. With that information, we split the enriched proteins of each condition in four subsets:1) proteins from the pull-down assay, 2) proteins from the pull-down exclusively enriched for this condition, 3) pull-down proteins that interact with RpoC in STRING and 4) pull-down proteins that do not interact with RpoC in STRING. We then performed a functional enrichment of each subset using DAVID (https://davidbioinformatics.nih.gov/). Data was cleaned from unspecific bindings via 1-not considering the proteins that were only captured under no-tagged strain (17 proteins) and 2-spotting where no-tag condition showed more abundance than the other conditions (108 proteins). Eventual cleaned data contains 307 proteins captured under all conditions.

### 2.6. *In silico* modelling

The 22 proteins that were enriched on every condition (see below) based on the criteria explained in the text were selected for further structural analyses. To this end, they were combined in dimers and modelled using AlphaFold2 (version 2.3.2, template date = 2023-05-14) to generate the ipTM and pTM score of the dimers to propose some bigger complexes later. RNA polymerase core (two β subunits, β and β’ subunits and ω subunit) was modelled with any enriched proteins on any condition using AlphaFold3 server (https://alphafoldserver.com/). Other complexes that could be of interest based on dimers scores were modelled with AlphaFold3 as well. Visualisation and alignment of complexes was done using Pymol software (https://www.pymol.org/)

## 3. RESULTS AND DISCUSSION

### *3*.*1* Tagging the C-terminus of the RpoC (β’) subunit

Because RNA polymerase (RNAP) performs one of the most essential functions in the cell, genetic modification of its subunits must be undertaken with caution at is touches the key actor of the gene expression flow. A potentially permissive insertion site was identified at the 3′ end of the *rpoC* gene, the structural modelling of which indicated that the corresponding region in the β′ subunit was located within an exposed protrusion on the surface of the RNAP holoenzyme, making it suitable for the introduction of affinity tags without, in principle, disrupting enzyme function (Fig. 1B). On this basis, 3 modifications were separately introduced immediately upstream of the native TAA stop codon. The detailed protocol to introduce them in the chromosomal *rpoC* gene is explained in Supplementary Fig. S1 and sketched in Fig. 1A. In sum, DNA sequences encoding a 10xHis tag, a c-Myc-tag, or an E-tag were added at the end of the ORF each for a different purpose. The poly-His peptide enables cost-effective purification of RNAP complexes, whereas the c-Myc and E-tags offer high specificity for immunoprecipitation and protein identification (Terpe, 2003). To improve tag accessibility and minimize potential steric hindrance, flexible Gly residues were inserted between the β′ subunit C-terminus and the corresponding label. Successful integration in the right genomic site was verified with PCR using primer pairs as indicated in Supplementary Table S1. Amplification products of the expected size (∼0.7 kb) confirmed correct chromosomal insertion and localization insert (Fig. 1C). To verify attachment of the tags to the β′ subunit of RNAP Western blot analysis of whole-cell extracts separately probed with anti-His, anti c-Myc and anti-E-tag antibodies demonstrated the presence of each of the epitopes in a protein band of ∼ 150 Kd in the gels. Given that the affinity tags have an approximate molecular mass of 1 kDa, their addition represents only a minor increase of the molecular mass of ∼ 155 kDa expected for RpoC. Note that the cognate in the Western blots appeared as well defined bands with no indication of proteolysis or instability, suggesting that this procedure labelled every single β′ in the cells. The resulting RNAP-tagged strains are called hereafter *P. putida* TAS108 (β′-10xHis), *P. putida* TAS110 (β′-Myc) and *P. putida* TAS114 (β′-E).

In order to verify whether insertion of the tags caused any gross perturbation on microbial physiology, we run growth curves in LB medium to compare their growth rates. As shown in Fig. 2A, doubling times of the RNAP-tagged bacteria were basically indistinguishable from that of the wild-type KT2440 strain. Similarly, visual inspection of colony development on agar plates (Fig. 2B) exposed no morphological differences among them after 3 days of growth on M9-citrate. Taken together, these results indicated that adding different tags to the C-terminus of RpoC had little or no influence on the overall physiology of *P. putida*, thereby enabling their use as baits for exploring the whole RNAP interactome. Among the tags entered in RpoC, c-Myc was selected for subsequent immunocapture experiments because its small size, high immunoreactivity and efficient recognition by well-characterized anti-Myc antibodies make it particularly suitable for immunoprecipitation and co-immunoprecipitation of native protein complexes (Hillman *et al*., 2001; Young *et al*., 2012; Zhao *et al*., 2013). The results discussed hereafter are thus based on studies done on strain TAS106 (*P. putida* KT2440 *rpoC*::MycTag *att*Tn7::PBG37-msfGFP; Supplementary Table S2)

**Figure 2.**
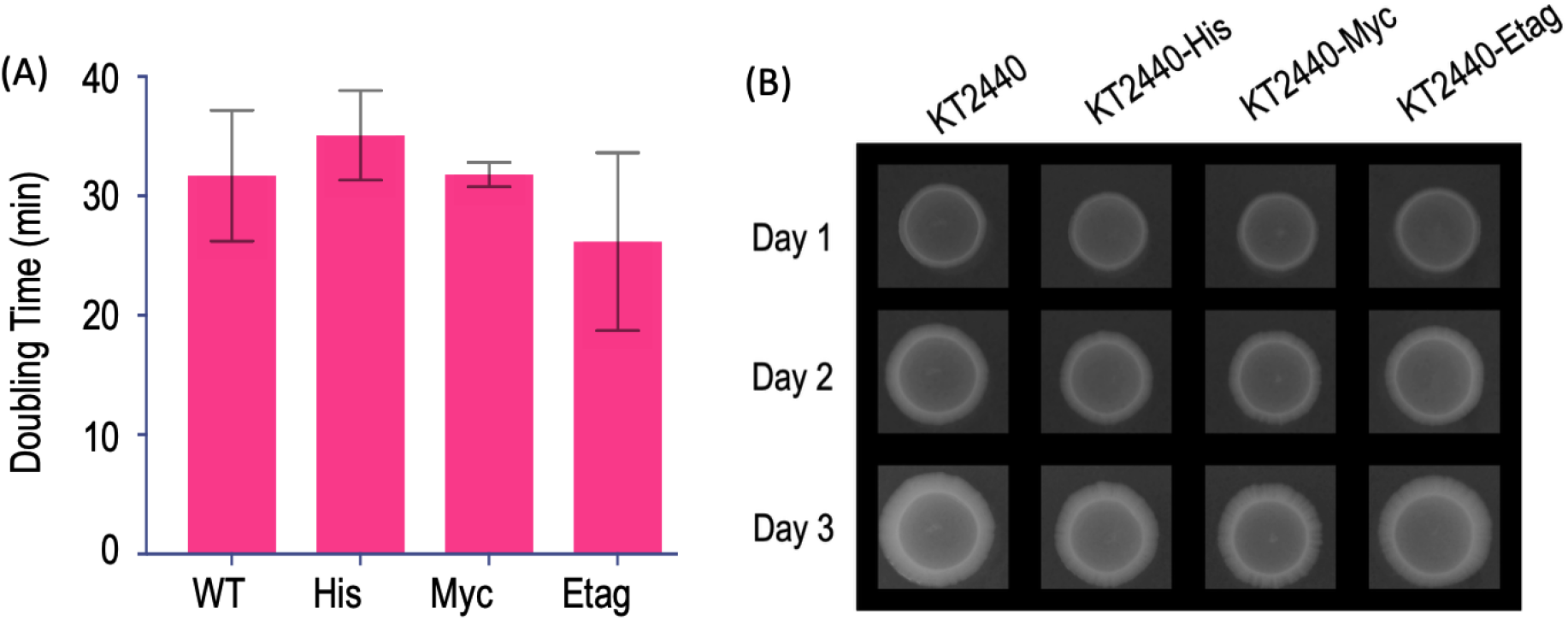
Gross phenotypic characteristics of tagged-RNAP *P. putida* strains. (A)Doubling times of each of the *P. putida* strains bearing tagged variants of RNAP strains. Each derivative was cultured in LB medium and their growth rates at mid-exponential phase calculated as shown. (B) Colony morphology. Each of the strains under study spotted on 1% agar in M9-citrate minimal medium and incubated for the time indicated. Note no conspicuous differences among the different variants

### 3.2 Rationale for sampling the molecular entourage of RNAP

Gene expression is arguably the central process that enables living cells to survive and adapt to changing environments. In bacteria, the decision to express a particular gene emerges from the integration of multiple regulatory layers that respond to a wide variety of signals, including physicochemical conditions, nutrient availability, and physiological cues reflecting the cell’s internal state. At the core of this regulatory network lies the transcriptional machinery, composed of RNA polymerase (RNAP), sigma factors, transcription factors, and a variety of termination and antitermination proteins that collectively determine when, where, and to what extent genes are transcribed (Browning and Busby, 2016; Ray-Soni *et al*., 2016). Although these components constitute a minimal machinery required for transcription *in vivo*, accumulating evidence suggests that bacterial RNAP operates within a much larger and more complex molecular environment. Computational analyses of gene co-expression and evolutionary conservation, together with experimental approaches such as immunoprecipitation and protein interaction studies, have revealed a growing number of proteins physically associated with the transcriptional apparatus (Murakami and Darst, 2003; Lee *et al*., 2008; Jin *et al*., 2013; Kohler *et al*., 2017; Kriel *et al*., 2020; Muskhelishvili *et al*., 2021). This view has led to the concept of transcription as a highly coordinated process carried out by a dynamic multiprotein complex that traverses the chromosome while simultaneously coupling mRNA synthesis to downstream cellular processes, including translation (Blaha and Wade, 2022).

A key unresolved question is whether the ensemble of proteins associated with RNAP remains largely constant across bacterial species and physiological states, or whether it undergoes continuous remodelling in response to environmental and cellular cues. In the latter scenario, the recruitment and release of accessory proteins could provide an additional layer of regulation, modulating transcriptional output according to the physiological context of the cell. To explore this possibility, we developed a protocol for the non-disruptive immunoprecipitation of proteins associated with the β′ subunit of RNAP in *P. putida* (see above). Importantly, the procedure was performed without chemical cross-linking, thereby facilitating the recovery of stable protein complexes under conditions that more closely reflect their native state. The protocol was applied to cells grown under distinct physiological conditions, including rapid growth in rich medium (exponential phase), stationary phase, and prolonged heat stress. While these conditions represent only a small subset of the many physiological states experienced by bacterial cells, they provide a useful framework for examining changes in the composition of the RNAP-associated protein network under contrasting environmental situations. The overall experimental workflow is summarized in Fig. 3. Briefly, proteins co-immunoprecipitating with the RNAP β′ subunit (RpoC) were compared across stationary-phase cultures (Condition S2), exponentially growing cells (Condition S3), and heat-shocked cultures (Condition S4). As a control, we analysed an otherwise identical *P. putida* strain lacking the chromosomal immunotag used for protein capture but still treated with the anti-Myc antibody (Condition S1). Following immunoprecipitation, protein complexes were isolated and analyzed by LC–MS/MS proteomics as explained in the Materials and Methods section. The resulting datasets were subsequently processed computationally, with particular attention given to the proteins exhibiting the most pronounced condition-dependent differences in abundance. Complete raw proteomic data are provided in Supplementary Table S3.

**Figure 3.**
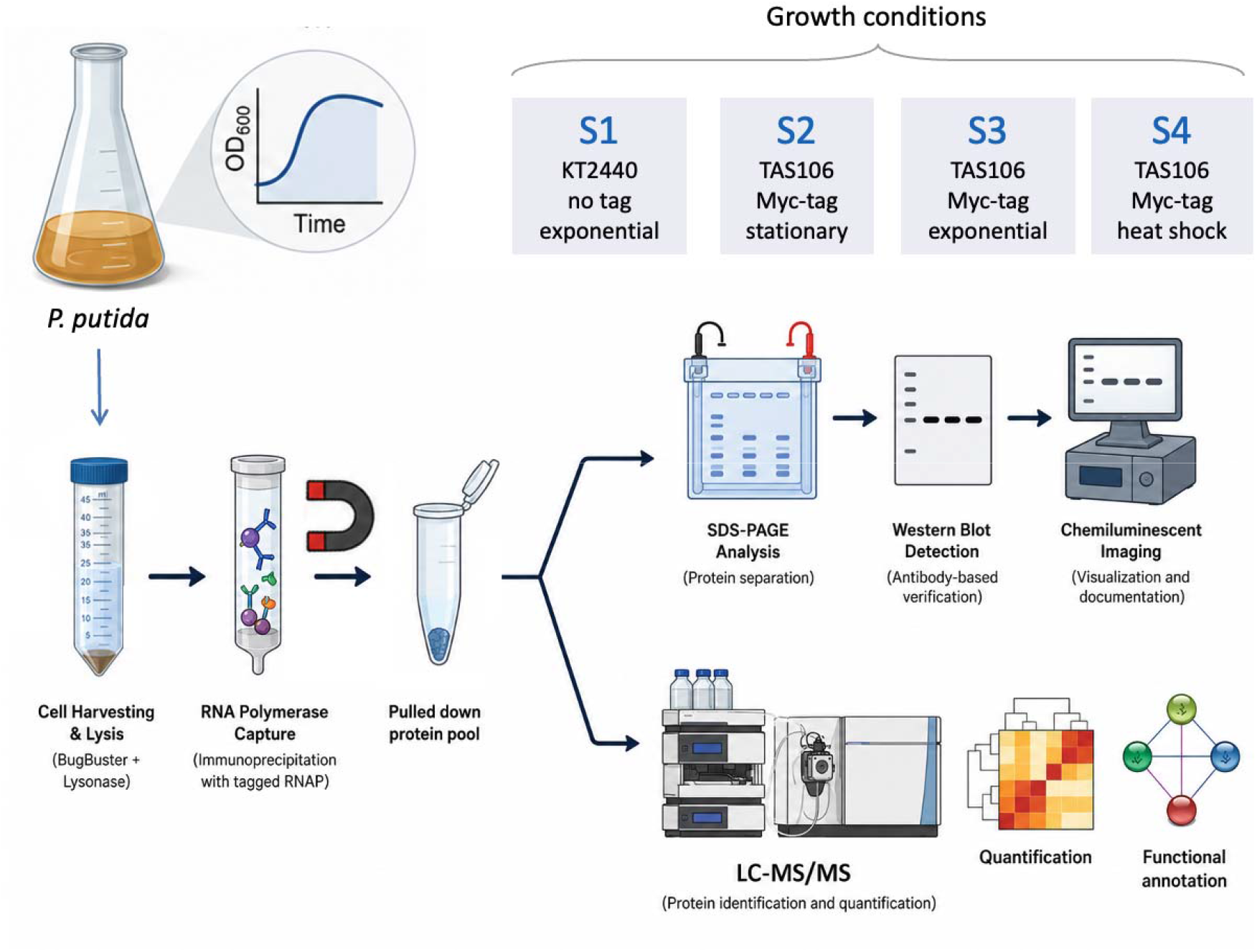
Workflow of pull down experiments for sampling the RNAP interactome of P. *putida*. Cultures were grown in aerated flasks using the medium and conditions described in the text and subjected to one of four experimental conditions (S1-S4). In all cases, cultures were inoculated from overnight cultures at an initial optical density OD_600_ of 0.02. For conditions S1, S3, and S4, cells were allowed to grow to an OD of 0.4 and were then either harvested directly or subjected to heat stress for 3 h, as specified for this condition. For condition S2, cultures were instead allowed to grow overnight. At the end of the corresponding growth or stress period, all cultures were normalized to an OD of 2.0 to ensure that equivalent numbers of cells were collected for subsequent analysis. Cells were then harvested and processed for immunomagnetic precipitation to isolate the protein fraction captured by the c-Myc tagged RNAP. The resulting samples were either analyzed by polyacrylamide gel electrophoresis to assess sample quality or submitted to the protein analysis platform for proteomic characterization.

### 3.3 The core interactome of P. putida RNAP

The immunoprecipitation experiments sketched in Fig. 3 generated proteins pools for each condition, out of which 307 species could be unequivocally identified and their level in each of the samples compared qualitatively and quantitatively. The criterium for considering a given protein associated to the bait RpoC-Myc was being overrepresented > 10 times in respect to the same protein in the reference control. The Venn diagram of Fig. 4 summarizes the result of this operation, indicating the groups of proteins pulled down with the β′ subunit of the polymerase in each situation. Note that there is a group of 22 proteins that appeared in all conditions and which can thus be considered the habitual molecular companion of the enzyme in vivo. These expectedly included all subunits of the core polymerase: RpoA (α subunit), RpoB (β) RpoC (β’) RpoZ (ω). Interestingly, the sigma factor habitually considered housekeeping (RpoD, σ^70^) was not shared in all conditions, while RpoH (heat shock σ^32^) and RpoN (σ^54^) showed up across all physiological states tested. The whole complement of Nus products (NusA, NusB, NusE/RpsJ, NusG) involved in mRNA elongation, termination, antitermination and transcription/translation coupling (Sen *et al*., 2008) were also associated to the RNAP in all conditions. This was somewhat expected in view of their functionality along the transcription process, which unavoidably required interactions with the RNAP core. In addition, the pool of proteins co-precipitated under all experimental conditions included RpsO (ribosomal protein S15, (Nikulin *et al*., 2000)), RlmL a 23S rRNA methyltransferase involved in large ribosomal subunit biogenesis (Lesnyak *et al*., 2006; Kimura *et al*., 2012), SuhB, a factor implicated in the maturation and assembly of the 30S ribosomal subunit (Singh *et al*., 2016), and DeaD (also known as CsdA, cold-shock DEAD-box protein A), an ATP-dependent RNA helicase involved in the remodelling of RNA structures and ribonucleoprotein complexes, with established roles in ribosome biogenesis, translation and RNA turnover (Iost and Dreyfus, 2006) (Jarmoskaite and Russell, 2014). Also related to nascent RNA management, the chaperone Hfq was consistently recovered in all RNAP pull-downs. This association is consistent with previous evidence that Hfq can interact with RNAP and, importantly, can be recruited co-transcriptionally to nascent RNAs as they emerge from the transcription complex, thereby positioning Hfq at an early interface between transcription and post-transcriptional regulation (Sukhodolets and Garges, 2003; Kambara *et al*., 2018; Rodgers *et al*., 2023). Furthermore, we found LepA (EF4), a highly conserved ribosome-dependent GTPase implicated in translational quality control, can promote reverse translocation (back-translocation) of the tRNA–mRNA complex and has been proposed to facilitate recovery of improperly translocated or stalled ribosomes, particularly under stress conditions (Qin *et al*., 2006; Pech *et al*., 2011; Yamamoto *et al*., 2014). In contrast, we failed to detect association of RNAP to any typical transcription factor, whether repressors or activators. Notably, DksA was not detected in any of the RNAP pull-downs, despite being a well-established RNAP-binding protein. DksA binds directly to the secondary channel of RNAP and acts synergistically with the alarmone ppGpp to modulate transcription, with a major ppGpp-binding site formed at the DksA–RNAP interface (Ross *et al*., 2016). It is plausible that both TFs and DksA behave as transient regulatory ligands of the polymerase rather than constitutive components of the holoenzyme and thus represent a very minor portion of the interactome at any given time.

**Figure 4.**
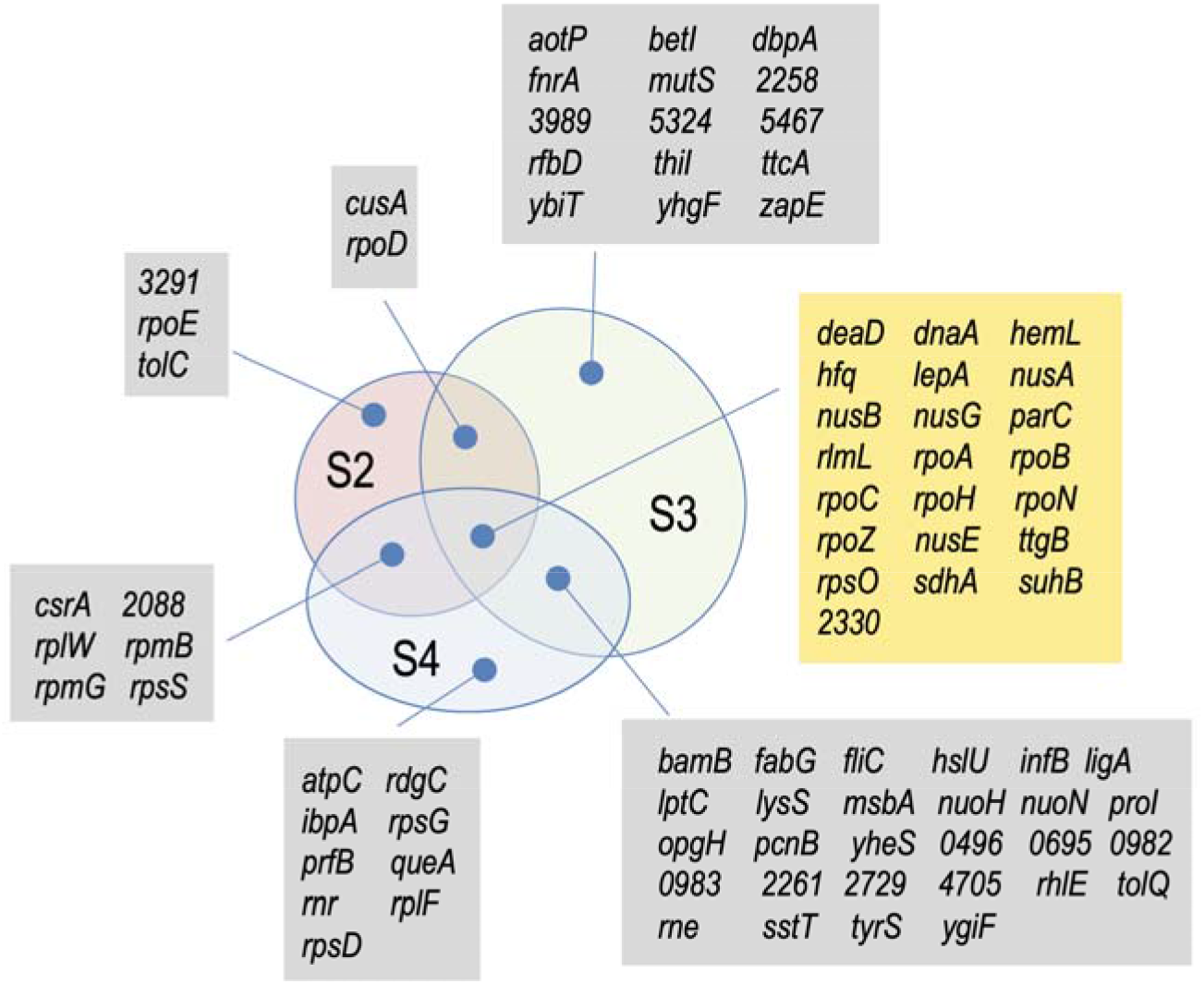
Condition-dependent composition of the RNA polymerase interactome in *Pseudomonas putida*. The Venn diagram shows the proteins unequivocally identified as co-immunoprecipitating with Myc-tagged RNA polymerase under the three physiological conditions analyzed (S2, S3, and S4), grouped according to their occurrence in individual or multiple conditions. The central overlap (yellow box) defines the basal RNA polymerase interactome, comprising 22 proteins that reproducibly co-precipitated with RNA polymerase under all three conditions and fulfilled the stringent selection criteria described in the Materials and Methods. This conserved set includes RNA polymerase subunits (RpoA, RpoB, RpoC, RpoH, RpoN, and RpoZ), the transcription-associated factors NusA, NusB, NusE, and NusG, as well as several proteins involved in RNA metabolism, ribosome function, chromosome maintenance, and cellular metabolism (see text for further discussion). Proteins detected exclusively under each physiological condition, as well as those shared between two conditions, are indicated in the corresponding sectors of the diagram. Although condition-specific and shared proteins are shown for comparison, only the 22 proteins common to all three conditions were retained for detailed analysis of the basal RNA polymerase interactome.

### 3.4 From interactome to expressome

Given the triplicate sampling procedure and the stringent criteria for selecting significant interactions, we argue that all of the above mentioned proteins in the pulldowns of all conditions were meaningful and likely to include much of what has been called the *expressome* of *P. putida* ie. the multi-protein molecular machines that run the gene expression flow within their native context (Kohler *et al*., 2017). Since the methods used average the interactome of the RNAP among cells and promoter strengths in individual cells, these associations point to formation of highly integrated gene-expression assemblies at given sites. For instance, the simultaneous detection of SuhB, NusA, NusG, NusB, ribosomal proteins (particularly NusE/S10), LepA, Hfq, rRNA-processing factors, and RNA helicases such as DeaD suggests that the immunoprecipitation had captured a molecular entity considerably larger than RNA polymerase alone in which transcription, rRNA maturation, and ribosome biogenesis, are physically and functionally interconnected, possibly in the proximity of rRNA genes. Such an interpretation is fully consistent with an emerging view of gene expression in *P. putida* in which RNA polymerase is not merely the enzyme responsible for RNA synthesis, but rather serves as the central hub of a dynamic and physiologically responsive multiprotein network that integrates multiple layers of gene-expression control (Yuste *et al*., 2006; Muskhelishvili *et al*., 2021; Blaha and Wade, 2022). The presence of factors involved in ribosome assembly and RNA processing alongside the core transcriptional machinery supports the notion that RNA polymerase operates within a broader expression complex, integrating transcription with downstream events required for the production of functional ribosomes and, ultimately, protein synthesis.

Along the line, DnaA, the highly conserved initiator of bacterial chromosome replication, was also recovered among the RNAP-associated proteins. DnaA binds and oligomerizes at the chromosomal origin, *oriC*, to promote origin opening and replisome assembly (Ozaki and Katayama, 2009). Its association with RNAP is intriguing given the long-established functional interplay between transcription and replication initiation at *oriC*, where local transcription can facilitate DnaA-dependent origin opening (Baker and Kornberg, 1988). But still, the most surprising proteins found in all conditions were two metabolic enzymes. One was HemL (glutamate-1-semialdehyde 2,1-aminomutase), an enzyme of the tetrapyrrole biosynthetic pathway and ultimately heme, for respiratory electron-transfer and numerous redox-active enzymes (Ilag *et al*., 1991; Heinemann *et al*., 2008; Layer *et al*., 2010). Its recurrent association with components of the core gene-expression machinery raises the possibility that transient interactions between HemL and RNA polymerase may contribute to mechanisms that coordinate metabolic status, heme biosynthesis, and respiratory capacity with global transcriptional programs. Under this scenario, HemL could represent a moonlighting protein that, beyond its canonical metabolic function, acts as a sensor or integrator of the cellular metabolic state. The same could be true also for the second metabolic enzyme found in the core transcriptional pulldown: SdhA, the FAD-containing catalytic flavoprotein subunit of succinate dehydrogenase (respiratory Complex II), which couples oxidation of succinate to fumarate in the TCA cycle with electron transfer to the respiratory quinone pool (McNeil and Fineran, 2013). This enzyme occupies a unique position in cellular metabolism by directly linking the tricarboxylic acid (TCA) cycle to the electron transport chain through the oxidation of succinate to fumarate and the transfer of electrons to the respiratory quinone pool. Its association with RNAP may therefore reflect the capture of physiological information related to the respiratory and energetic state of the cell, suggesting a potential connection between transcriptional activity and central metabolic processes. TtgB was also identified among the proteins associated with RNAP. TtgB constitutes the inner-membrane transporter component of the tripartite RND-family TtgABC efflux pump, which contributes to the extrusion of organic solvents and other toxic compounds and thereby plays an important role in environmental stress tolerance (Segura *et al*., 1999; Ramos *et al*., 2002). Although a direct physical interaction between TtgB and RNA polymerase would not be expected, its recurrent detection may reflect a transient functional link between transcriptional activity and cellular stress-response mechanisms. The role of PP2330, also found in the mix could not be assigned.

### 3.5 Pairwise interactions among proteins of the core *P. putida* interactome

Having the list of enriched proteins of the core interactome, we checked at STRING database (https://string-db.org/, version 11.5) which of these proteins had a described interaction with RpoC with a high confidence score (< 0.900) and then performed analysis of the thereby predicted interactions using DAVID (https://davidbioinformatics.nih.gov/). In addition, the 22 proteins of the core interactome were combined pairwise in dimers and modelled using AlphaFold2 (version 2.3.2, template date = 2023-05-14) to generate the ipTM (interface predicted template modeling) and pTM (predicted template modeling) scores of potential interacting surfaces that could assist formation of larger complexes (these parameters are confidence metrics used by AlphaFold to evaluate the quality of predicted multi-protein complexes and interactions). The result of such a double-criteria (ie STRING-based interaction prediction and pairwise structural docking among all proteins of the core interactome) is summarized in Fig. 5. Expectedly a first group of interactions involves the components of the core RNAP (σ, β, β’ and ω subunits along with the most prevailing sigma factor (RpoH). These appear along with the whole set of transcription elongation factors (NusA, NusG, NusB, NusE) that couple transcription, translation, RNA folding, termination, and ribosome biogenesis, thereby signalling that our RNAP pull-downs did capture stable elongation complexes. As indicated above, the remaining factor SuhB appears also in the pulldown, but not in the pairwise analysis. On the other hand, RlmL and NusE/RpsJ makes a connection to ribosomal assembly and functioning. Specifically, NusE is also the ribosomal protein S10 which links transcription and translation in bacteria by first binding the 16S rRNA to ensuring accurate decoding during translation, after which it leaves the ribosome and joins the Nus transcription antitermination complex (Burmann *et al*., 2010). Yet, what is most remarkable in the sketch of Fig. 5 is the apparent central role of SdhA to nucleate many of the interactions of the complex. The recurrent appearance in all the pulldowns along with the STRING and AF2 predictions of interactions with other members of the basic interactome make unlikely trivial explanations of its presence and suggest instead an authentic participation in the complex. Metabolic enzymes are not infrequent in RNAP interactome studies (Klein *et al*., 2023; Stenum *et al*., 2023) although no specific mechanism of their role in such complexes have been clarified. One attractive scenario is that bacterial RNAP exists *in vivo* as the core of a larger supercomplex that transiently recruits RNA-processing factors, ribosomal proteins, translation factors, and selected metabolic enzymes that coordinate gene expression with metabolic state. This possibility does deserve further studies.

**Figure 5.**
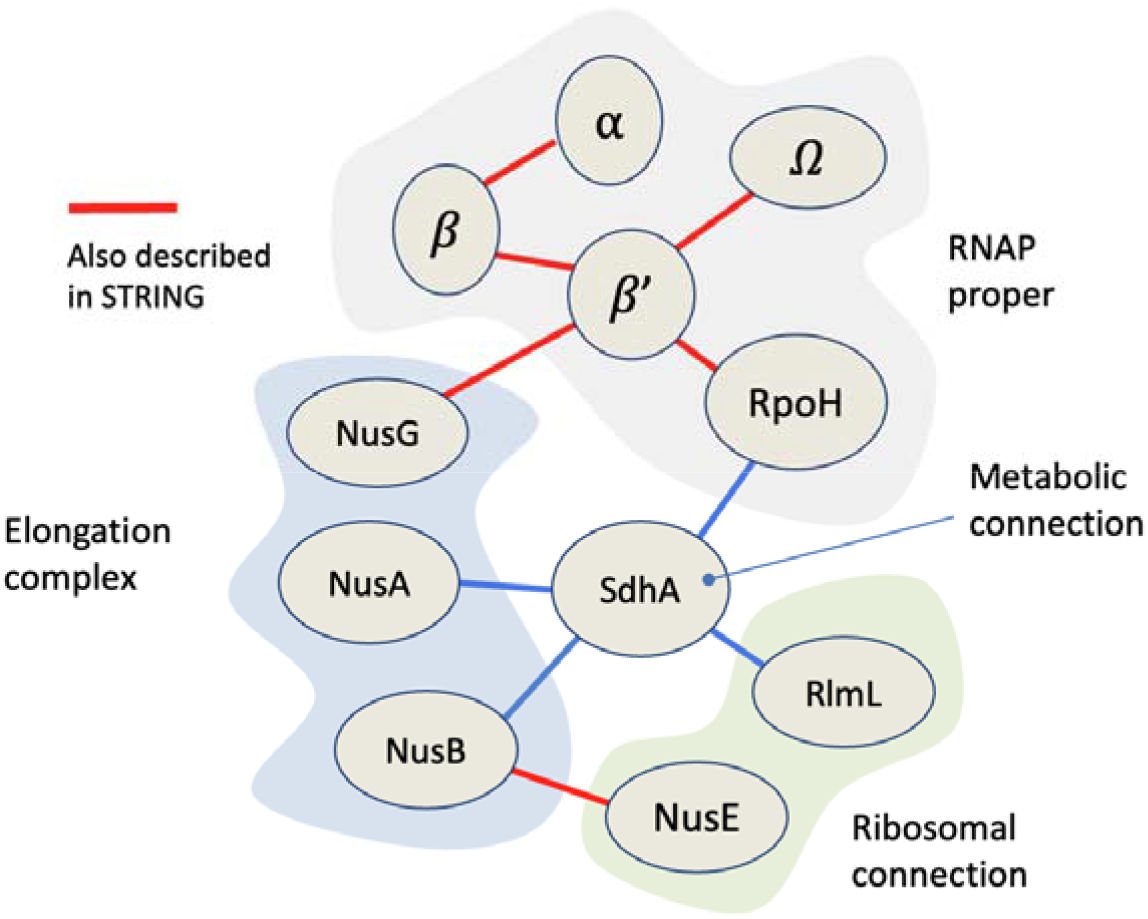
Interaction network within the basal RNA polymerase interactome *of Pseudomonas putida*. Pairwise structural interactions among components of the constitutive RNAP interactome of *P. putida*. Out of the 22 proteins shared in all of the pulldown pools, the 11 products shown in this scheme return high scores of surface-surface contacts between the partners indicated as predicted with AF2. In addition, some of them are anticipated with STRING to have structural and/or functional relation. The figure groups the detected proteins into functional modules—RNAP proper, elongation complex, ribosomal connection, and metabolic connection—while showing specific predicted or previously described interactions among them.

### 3.6 Compositional changes of the *P. putida* expressome across conditions

Fig. 5 not only defines the main RNA polymerase interactome but also reveals its remarkable compositional plasticity. Across the experimental conditions examined, a conserved set 21 proteins consistently co-precipitates with RpoC, representing—as argued above—the basic transcriptional machinery. Superimposed on this core, however, are condition-specific proteins whose association varies markedly with the physiological state of the cells. In some cases, these changes involve only a few proteins, whereas in others they are extensive. For example, conditions S3 and S4 share a large subset of additional interacting proteins, indicating that related physiological states recruit similar functional modules to the transcriptional apparatus. Because our analysis applied stringent criteria for defining bona fide interactors, these differences are unlikely to arise from experimental noise alone and instead point to genuine remodelling of the RNAP interactome in response to environmental conditions.

Although it is inevitable that the methodology fails to capture some true interactors while allowing a number of possible contaminants to remain, these limitations do not obscure the overall pattern. Instead, the data consistently demonstrate that the protein environment surrounding RNA polymerase is dynamic, with distinct constellations of associated proteins acting under different physiological conditions. Note that many of the condition-specific interactors are directly involved in gene expression, supporting the notion that transcription is coordinated with RNA processing, translation, and other gene-expression-related processes. Others have no obvious connection to transcription, suggesting that the observed complexes may reflect multiple layers of protein interactions. In this view, RNA polymerase associates not only with its immediate functional partners but also with proteins connected through secondary or tertiary interaction networks (Fig. 6), thereby linking transcription to broader aspects of cellular physiology. Although the present data do not allow the precise functional contribution of each individual interactor to be established, the observed remodelling of the interactome strongly suggests that RNA polymerase functions as more than a catalytic machine for RNA synthesis. Instead, it appears to act as a dynamic molecular hub that integrates physiological signals and coordinates transcription with the changing metabolic and regulatory state of the cell.

**Figure 6.**
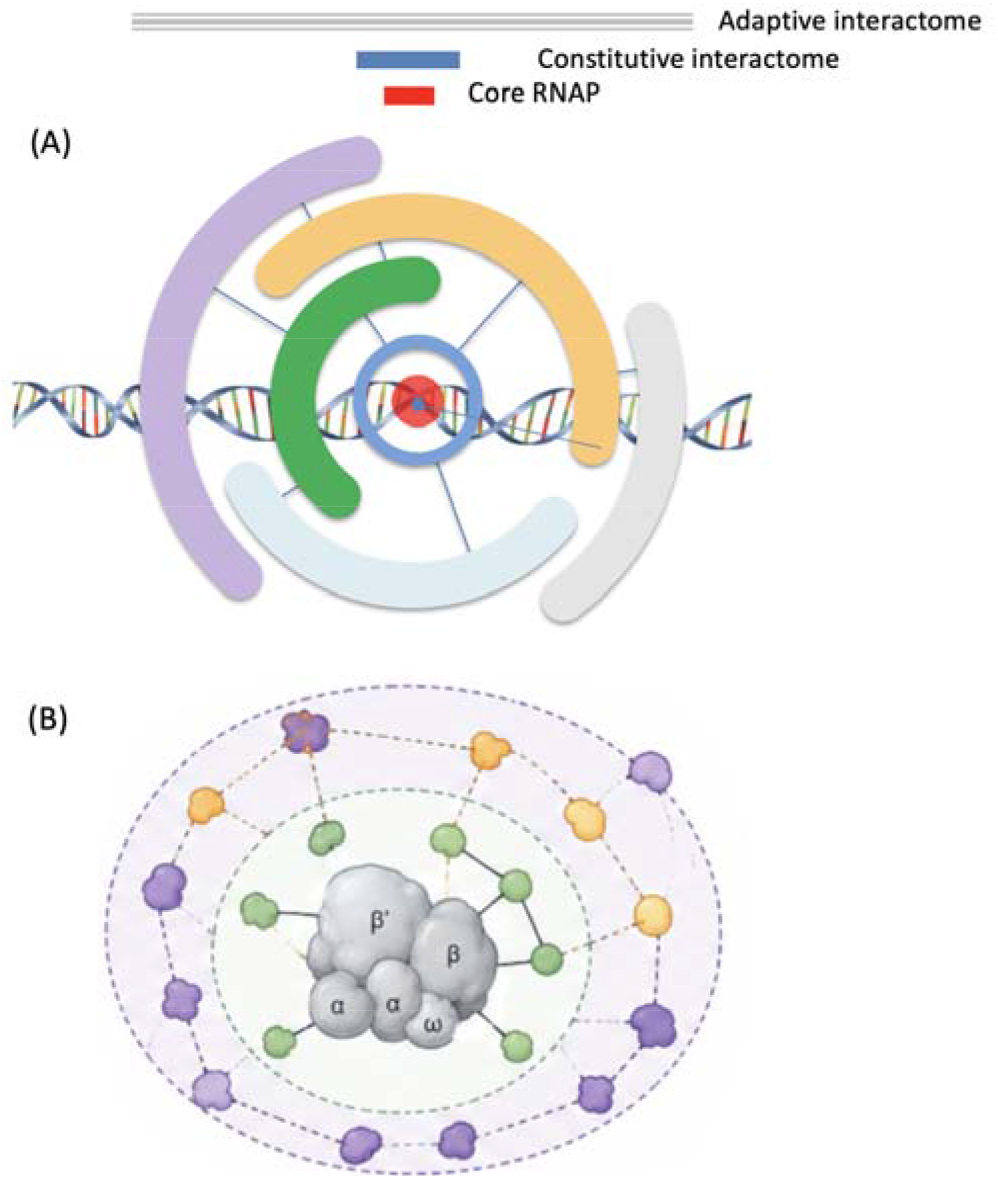
A core-and-shell model of the condition-dependent RNA polymerase interactome. Schematic representation of the proposed organization of the *Pseudomonas putida* RNAP interactome as a stable core surrounded by progressively more condition-dependent protein associations. **(A)** The RNAP core (red) is associated with a constitutive set of proteins (blue) that defines the basal interactome maintained across the physiological conditions examined. Beyond this conserved configuration, additional proteins may associate directly or indirectly with RNAP in a condition-dependent manner, generating an adaptive interactome whose composition varies with the physiological state of the cells. The different outer sectors schematically represent alternative sets of proteins recruited under different growth or stress conditions and are not intended to indicate fixed structural arrangements. **(B)** Alternative representation of the same concept as concentric interaction layers. RNAP and its constitutively associated proteins form the innermost, conserved interaction network, whereas additional layers represent increasingly indirect and/or condition-specific associations. Thus, rather than constituting a single invariant molecular complex, the RNAP interactome can be visualized as a persistent basal scaffold embedded within a larger and dynamically reconfigurable network of protein interactions that responds to the physiological state of the cell.

This notion was further strengthened by inspecting the actual variation of the main participants of the interactome in each of the conditions tested. To this end, we just focused on the top over-represented 15 proteins (including RNAP core subunits) based on the normalized abundances of each of them under the specific physiological states. Fig 7 shows the combined bar plot representation of their relative abundance. The stability of the 3 main components of RNAP (RpoA, B and C) serve as an internal control for the variations of the others, some of which changed drastically. Along the line, we also inspected separately the changes in the sigma factor pool in each of the conditions. Sigma factors are expected to be among the most dynamic components of the RNA polymerase complex. Unlike the core enzyme (α_2_ββ′ω), sigma factors exchange rapidly in response to environmental cues. As shown in Fig. 8., while RpoH and RpoN appeared in all pulldowns, the majoritary pool of sigmas in stationary phase involved RpoS and RpoE, what makes a lot of sense in cells under starvation and possible oxidative stress. Thus, finding distinct sigma factor profiles under S2-S4 conditions supports that the pull-downs capture physiologically distinct transcriptional states and that transcription machinery itself is being reconfigured in response to environmental cues.

**Figure 7.**
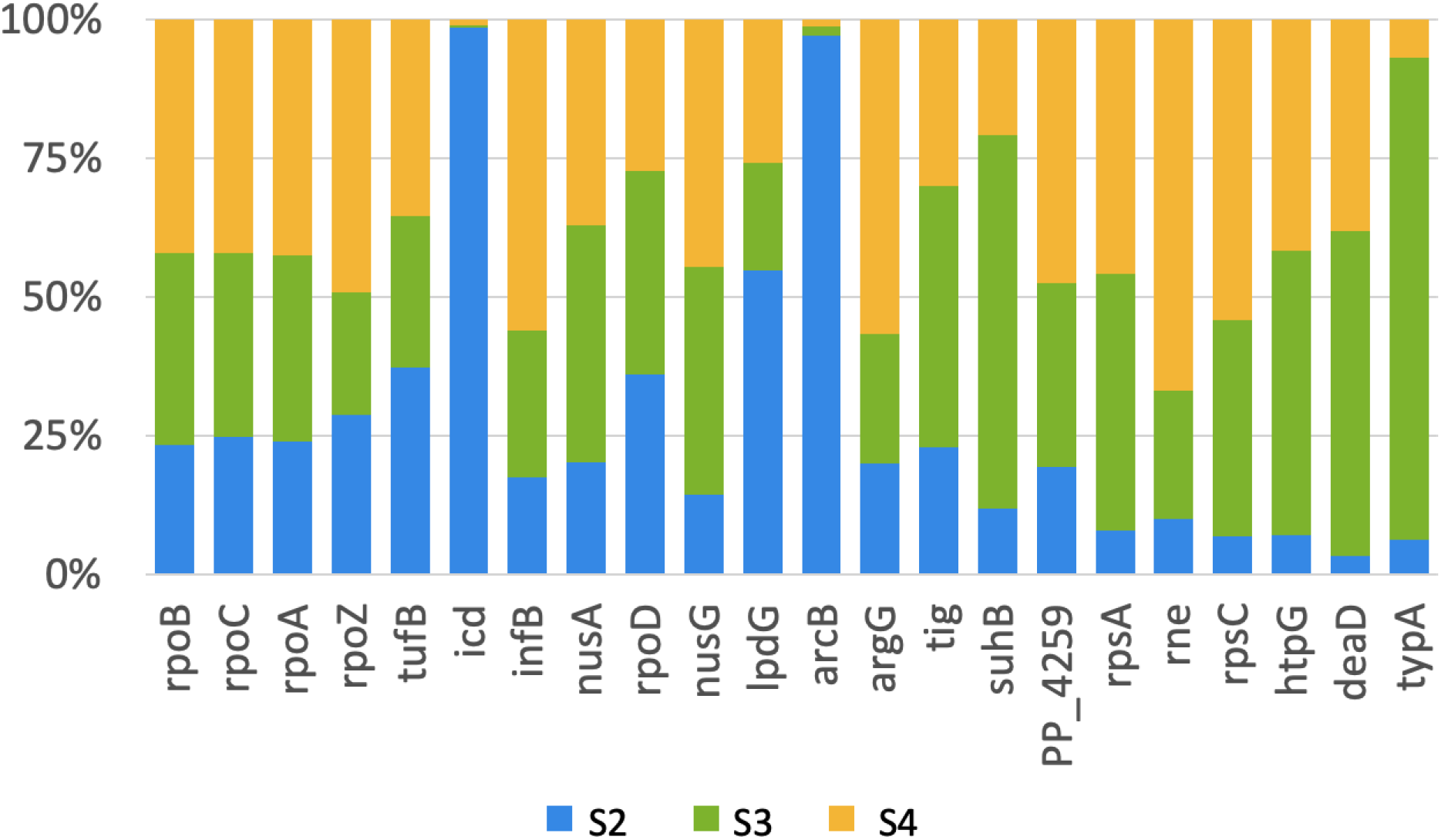
Remodeling of the RNAP interactome under different physiological regimes. Relative distribution of proteins associated with the *Pseudomonas putida* RNA polymerase (RNAP) interactome under three physiological conditions: exponential growth (S2, blue), stationary/ovemight growth (S3, green), and heat shock (S4, yellow). The graph includes 18 selected proteins of the RNAP-associated protein pool, apart of the four core RNAP subunits, and illustrates how their representation within the interactome shifts according to the physiological state of the cells. For each protein, normalized abundance values obtained under the three conditions were scaled to a total of 100%, so that each stacked bar depicts the relative contribution of S2, S3, and S4 to the overall signal detected forthat protein. Thus, the graph is intended to visualize changes in the relative association of individual proteins with RNAP across physiological states, rather than differences in their absolute cellular abundance. Some proteins show preferential association with RNAP during exponential growth, whereas others become comparatively enriched during stationary phase or following heat shock. Together, these changes illustrate the dynamic nature of the RNAP interactome: a conserved basal configuration is maintained while the relative representation of associated proteins is remodelled in response to the

**Figure 8.**
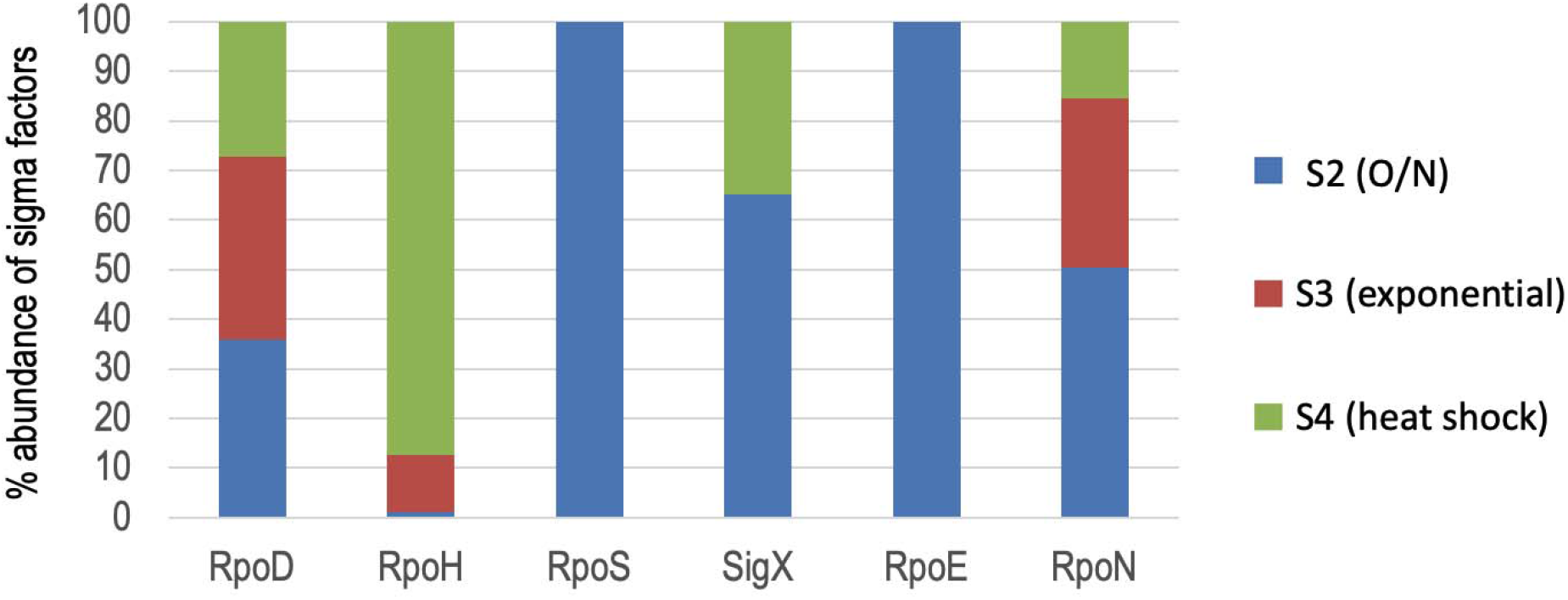
Condition-dependent association of sigma factors with RNA physiological state of the cell. Relative association of sigma factors with *P. putida* RNAP under three physiological conditions: S2 (overnight/stationary-phase culture; blue), S3 (exponential growth; red), and S4 (heat shock; green). Abundance values obtained for each sigma factor under S2-S4 were first normalized against the corresponding signal detected in the S1 control condition. To avoid division by zero when no signal was detected in S1, a negligible value (10^−^□) was added to the denominator. The resulting normalized values were then expressed, for each sigma factor, as percentages of the sum of its normalized abundance across S2, S3, and S4. Accordingly, each stacked bar sums to 100% and represents the relative distribution of a given sigma factor among the three physiological conditions, rather than its absolute abundance or a comparison of abundance between different sigma factors. The analysis reveals marked condition-dependent differences in sigma-factor association with RNAP. Whereas some factors are recovered under several physiological regimes, others show a strong preference for a particular condition. For example, RpoS and RpoE were detected exclusively under S2 among the three conditions compared and therefore appear as 100% S2 in the normalized representation. Overall, the data illustrate the physiological plasticity of RNAP composition through differential recruitment of alternative sigma factors.

### 3.7. Conclusion

To our knowledge, this work provides a first condition-dependent view of the *P. putida* RNA polymerase interactome *in vivo*. By introducing a minimally invasive chromosomal tag into the β′ subunit of RNAP, we established a robust strategy for isolating native transcriptional complexes under different physiological conditions. Beyond recovering the expected core transcription machinery, our analyses revealed a conserved set of proteins linking RNAP to RNA maturation, ribosome assembly and translation, together with a broader collection of condition-specific partners whose composition changes according to the physiological state of the cell. As a matter of fact, the RNAP interactome appears to comprise proteins with very different residence times in the complex. Some, such as NusA and NusG, behave as relatively stable members of the transcription complex, whereas others, like DksA—let alone typical transcription factors—are transient regulators that may escape detection with our approach despite playing major physiological roles. In contrast, the recovery of unexpected proteins under specific conditions indicates the formation of higher-order, condition-dependent molecular assemblies that extend beyond the canonical transcription machinery. These observations support the emerging view that bacterial RNA polymerase functions not as an isolated enzyme but as the central hub of a dynamic expressome that coordinates transcription with multiple aspects of cellular physiology. The unexpected association of metabolic enzymes with RNAP further suggests that metabolic status may be communicated directly to the transcriptional apparatus through higher-order molecular assemblies. Extending this approach to a wider range of environmental conditions, genetic backgrounds and perturbations should help define the architecture, dynamics and functional significance of these complexes, providing an additional regulatory layer for understanding bacterial gene expression and adaptation.

## Author Contributions

The study was designed by VdL, HT and FP. *in vitro* and *in vivo* experiments were run by HT. *in silico* analyses were run by BP. All Authors contributed to the data handling and bioinformatic analyses. VdL and HT wrote the paper.

## Acknowledgements

We thank the Proteomics core facility of the CNB-CSIC for sample processing and analyses. No AI tools were used to create, alter, or manipulate original research data or results. All experiments were conducted at CNB-CSIC. This work was funded by the EMPOWERPUTIDA (H2020-LEIT-BIO-2014-1) and.

NYMPHE (CL6-2021-UE 101060625) Contracts of the European Union.

## Declaration of Generative AI and AI-assisted technologies in the writing process

During the preparation of this work, the authors used ChatGPT 5.5 in a limited manner to improve the readability and language of the manuscript. After using this tool, the authors reviewed and edited the content as needed and take full responsibility for the content of the publication.

## Ethics statement

This work does not involve any ethical issue

## Conflicts of Interest

The authors declare no conflicts of interest.

## Data Availability Statement

The data that supports the findings of this study are available in the supplementary material of this article.

## Strain Availability Statement

The engineered strains from this study are available upon request from Víctor de Lorenzo and Huseyin Tas.

## SUPPORTING INFORMATION

Additional supporting information can be found online in the Supporting Information section:

**Supplementary Figure S1**. Insertion of C-terminal immune-tags in the β’ subunit (RpoC) of RNAP.

**Supplementary Table S1**. List of primers used in this study (XLSX).

**Supplementary Table S2**. Strains used in this study.

**Supplementary Table S3**. Raw proteomic data.

---

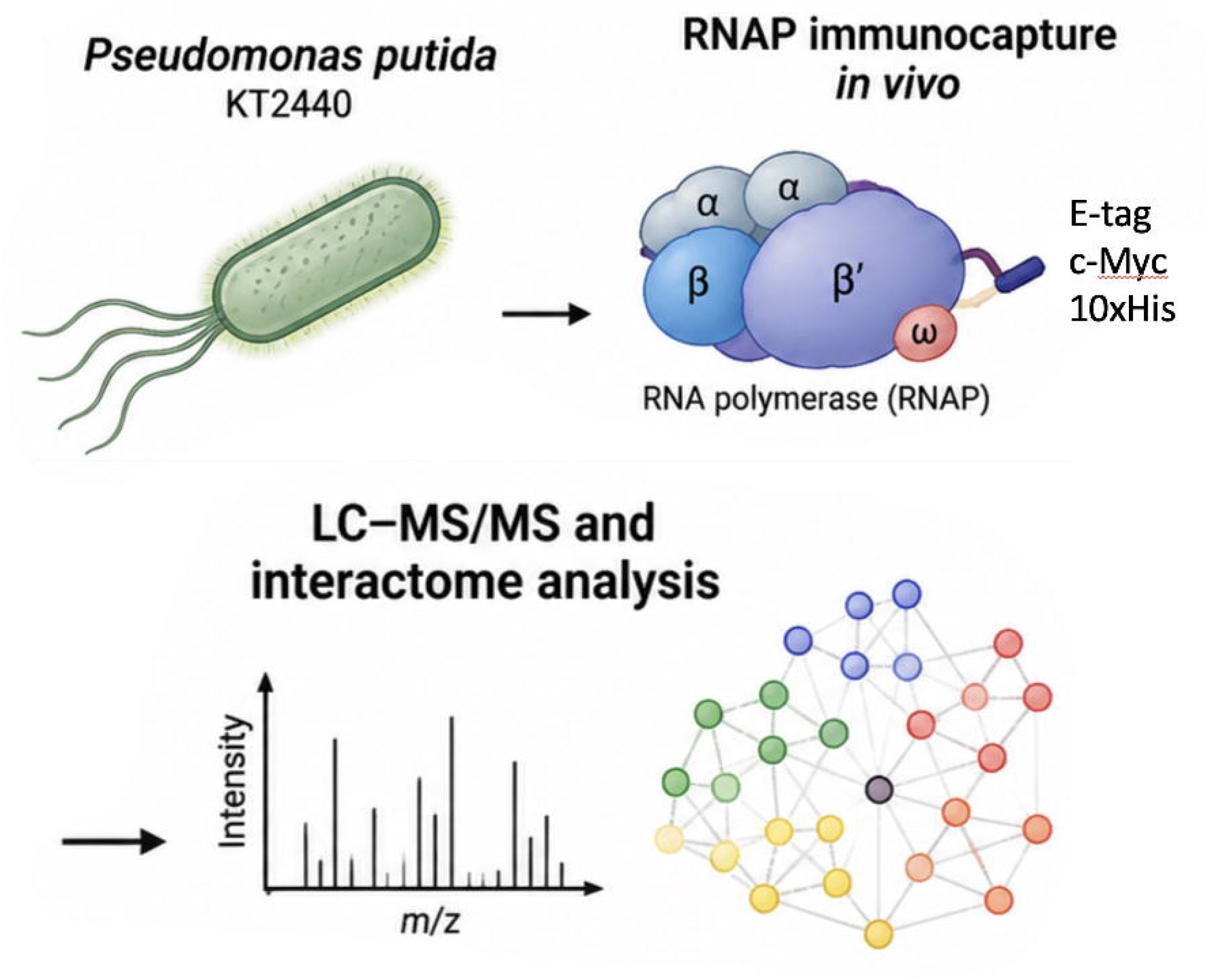

## Notes

### Competing Interest Statement

The authors have declared no competing interest.

