## Supplementary figure S1 for "The dynamic expressome of *Pseudomonas putida*: condition-dependent remodelling of the RNA polymerase interactome"

**Supplementary Figure S1.** Construction and verification of C-terminally tagged RpoC variants.

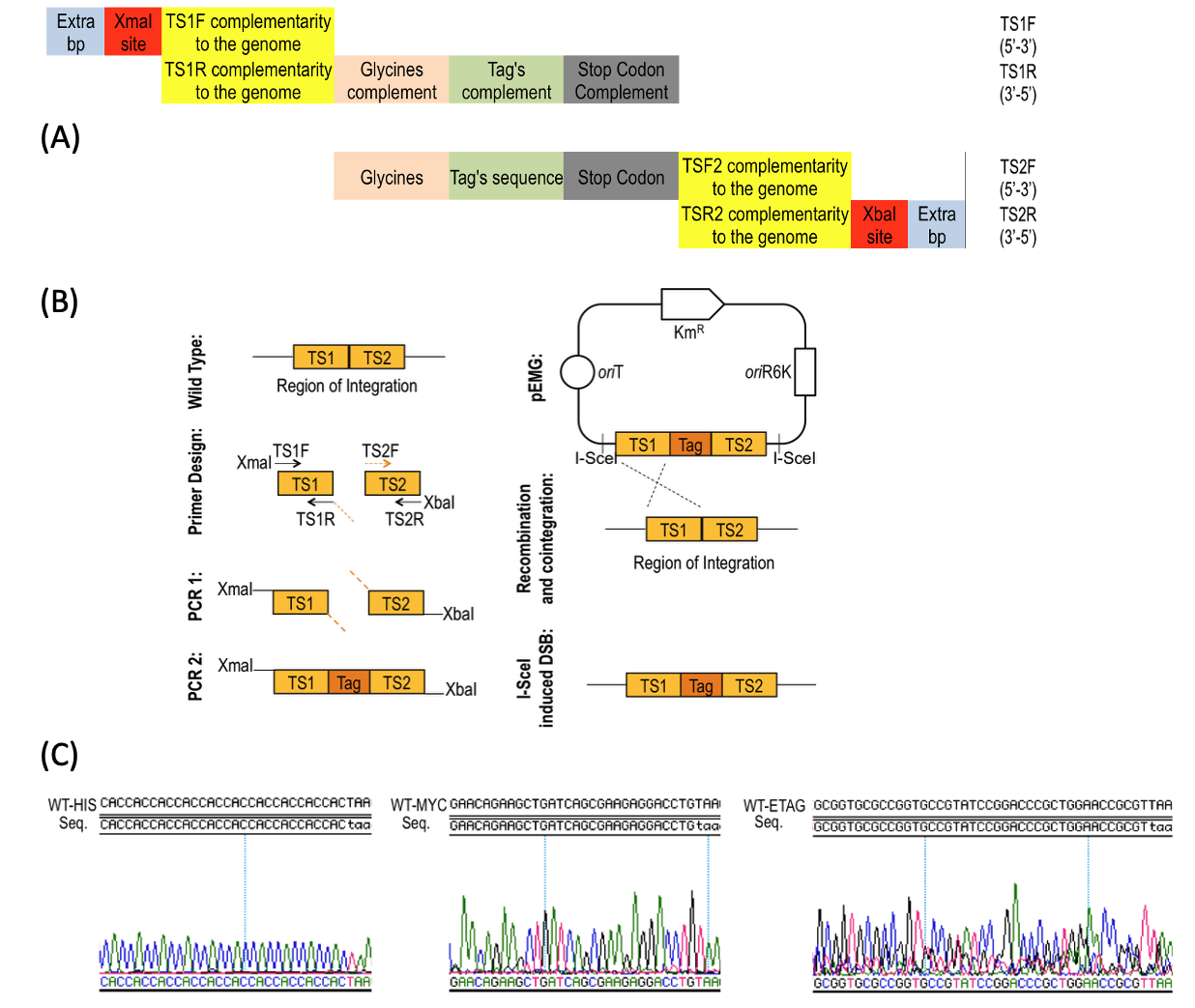

Schematic overview of the strategy used to introduce epitope tags at the C terminus of the RNA polymerase β′ subunit (RpoC). **(A)** Architecture of the primers used to generate the two targeting sequences (TS1 and TS2) flanking the *rpoC* integration site. Primers contained regions complementary to the chromosome together with additional sequences required for assembly, including the tag sequence, glycine-encoding linker, stop codon, XmaI restriction sites, and additional flanking nucleotides. The corresponding primer sequences are provided in Supplementary Table S1. **(B)** Workflow for construction and chromosomal integration of the tagging cassette. TS1 and TS2, corresponding to the genomic regions immediately flanking the intended insertion site, were first amplified by PCR. A second PCR assembled the two targeting sequences with the sequence encoding the desired C-terminal tag, generating the final TS1–Tag–TS2 fragment. The resulting cassette was cloned into the suicide vector pEMG and introduced into *P. putida*, where homologous recombination promoted cointegration of the plasmid at the *rpoC* locus. Subsequent I-SceI-mediated double-strand cleavage stimulated resolution of the cointegrate by a second homologous recombination event, yielding the desired chromosomal insertion of the tag while eliminating the plasmid backbone, according to the strategy described by (Martínez-García and de Lorenzo, 2011). **(C)** Verification of the engineered *rpoC* alleles by Sanger sequencing. Representative chromatograms are shown for the C-terminal His-, Myc-, and E-tagged variants, confirming the expected tag sequences and their correct insertion at the *rpoC* locus.
